# Hepatocytes deficient in nuclear envelope protein lamina-associated polypeptide 1 are an ideal mammalian system to study intranuclear lipid droplets

**DOI:** 10.1101/2022.06.27.497855

**Authors:** Cecilia Östlund, Antonio Hernandez-Ono, Samantha J. Turk, William T. Dauer, Henry N. Ginsberg, Howard J. Worman, Ji-Yeon Shin

**Affiliations:** Department of Medicine, Vagelos College of Physicians and Surgeons, Columbia University, New York, NY, USA; Department of Pathology and Cell Biology, Vagelos College of Physicians and Surgeons, Columbia University, New York, NY, USA; Peter O’Donnell Jr. Brain Institute, University of Texas Southwestern Medical Center, Dallas, TX, USA; Department of Neurology, University of Texas Southwestern Medical Center, Dallas, TX, USA; Department of Neuroscience, University of Texas Southwestern Medical Center, Dallas, TX, USA

**Keywords:** animal models, cell biology, lipid droplet, lamin, liver, mouse genetics, nucleus, high-fat diet, type 1 nucleoplasmic reticula, nutritional state

## Abstract

Lipid droplets (LDs) are generally considered to be synthesized in the ER and utilized in the cytoplasm. However, LDs have been observed inside nuclei in some cells, although recent research on nuclear LDs has focused on cultured cell lines. To better understand nuclear LDs that occur *in vivo*, here we examined LDs in primary hepatocytes from mice following depletion of the nuclear envelope protein lamina-associated polypeptide 1 (LAP1). Microscopic image analysis showed that LAP1-depleted hepatocytes contain frequent nuclear LDs, which differ from cytoplasmic LDs in their associated proteins. We found type 1 nucleoplasmic reticula, which are invaginations of the inner nuclear membrane, are often associated with nuclear LDs in these hepatocytes. Furthermore, *in vivo* depletion of the nuclear envelope proteins lamin A and C from mouse hepatocytes led to severely abnormal nuclear morphology, but significantly fewer nuclear LDs than were observed upon depletion of LAP1. In addition, we show both high fat diet feeding and fasting of mice increased cytoplasmic lipids in LAP1-depleted hepatocytes, but reduced nuclear LDs, demonstrating a relationship of LD formation with nutritional state. Finally, depletion of microsomal triglyceride transfer protein did not change the frequency of nuclear LDs in LAP1-depleted hepatocytes, suggesting that it is not necessary for the biogenesis of nuclear LDs in these cells. Together, these data show that LAP1-depleted hepatocytes represent an ideal mammalian system to investigate the biogenesis of nuclear LDs and their partitioning between the nucleus and cytoplasm in response to changes in nutritional state and cellular metabolism *in vivo*.

## INTRODUCTION

The lipid droplet (LD) is a dynamic cellular organelle composed of a core of triacylglycerols and sterol esters, a phospholipid monolayer and associated proteins. LDs are involved in multiple essential cellular functions including energy and lipid homeostasis (1, 2). Dysregulation of LD biogenesis and utilization has been connected to human diseases such as obesity, diabetes mellitus, non-alcoholic steatohepatitis, atherosclerosis, lipodystrophy, and neutral lipid storage disease (3). LDs have traditionally been considered to be synthesized and utilized only in the ER and cytoplasm. However, an electron microscopic study performed more than 60 years ago showed invaginations of the nuclear envelope containing LDs inside nuclei of mouse hepatomas and enlarged hepatocytes from mice fed a diet containing bentonite (4). Subsequent research identified a distinct endonuclear lipid domain in nuclei of normal rat hepatocytes (5, 6) and an electron microscopic study of human liver tissue from a patient with hepatitis C virus infection demonstrated a small percentage of hepatocytes with nuclear LDs (7). Feeding mice perfluorooctanoic acid also leads to accumulation of nuclear LDs in hepatocytes (8).

Research in cultured mammalian cell lines and non-mammalian model organisms has further addressed the biogenesis of nuclear LDs and their roles in metabolism. Nuclear LDs are associated with promyelocytic leukemia (PML) nuclear bodies in U2OS osteosarcoma and hepatocyte-derived cell lines (9, 10). When these cells are cultured in media containing oleic acid (OA), nuclear LDs associate with CTP:phosphocholine cytidylyltransferase ⍺ (CCT⍺), the rate-limiting enzyme in the CDP-choline pathway (9–11). CCT⍺ alternates between an inactive nucleoplasmic form and a nuclear membrane-bound active form (12, 13). In 3T3-L1 cells, CCT⍺ redistributes from the nucleoplasm to the nuclear envelope and cytosol, but does not colocalize with cytoplasmic LDs when OA is added to the culture media (14, 15). Nuclear LDs are rich in phosphatidic acid phosphatase and diacylglycerol, further suggesting that they are involved in lipid metabolism (10). Soltysik and colleagues (16) showed that nuclear LDs in hepatocyte-derived cell lines arise from apoB-free VLDL precursors in the ER lumen. These LDs enter into the nucleoplasm through type 1 nucleoplasmic reticula (11), which are invaginations of the inner nuclear membrane without outer nuclear membrane or nuclear pore complexes (17). However, U2OS cells lacking VLDL secretion machinery but containing CDP-choline and triacylglycerol synthesis enzymes in the inner nuclear membrane can generate nuclear LDs (10). Experiments in *Saccharomyces cerevisiae* show that the inner nuclear membrane is metabolically active and generates nuclear LDs (18, 19) and that increased unsaturated fat leads to a shift from nuclear LD formation to cytoplasmic synthesis at the outer nuclear membrane/ER (20). In *Caenorhabditis elegans*, nuclear LDs occur normally in intestinal and germ cells, and they are likely to be associated with nuclear damage only in the intestine (21). These experiments utilizing cultured cells and model organisms have been very informative. However, they are of limited generalizability as cultured cell lines may not be fully differentiated and the nuclear envelopes of model organisms, particularly in yeast that undergo a closed mitosis and do not have a nuclear lamina, differ from those in mammalian cells. The relationship between nuclear LDs and the nuclear envelope in cells of intact mammalian tissues may therefore be different.

In metazoans, the nuclear envelope consists of the inner and outer nuclear membranes, nuclear pore complexes, and the nuclear lamina. Most differentiated mammalian cells express A-type and B-type lamins, which are intermediate filament proteins that polymerize to form the lamina (22–25). Nuclear lamins also bind to integral proteins of the inner nuclear membrane and these interactions help maintain the lamina at the nuclear periphery. The outer nuclear membrane is directly continuous and shares most of its proteins with the ER, the major site of lipid biogenesis. While genetic research in humans has linked mutations in the gene encoding A-type lamins to partial lipodystrophy syndromes that primarily affect adipocytes (26–28), the role of the nuclear envelope in regulating cellular lipid homeostasis is poorly understood. Furthermore, the mechanism responsible for the generation of nuclear LDs in differentiated mammalian cells is largely unknown (29–31).

Our group has demonstrated that *in vivo* hepatocyte-specific depletion of lamina-associated polypeptide 1 (LAP1), an integral inner nuclear membrane protein, reduces VLDL secretion and causes accumulation of LDs within the nucleus and cytoplasm of mouse hepatocytes (32). One function of LAP1 is to bind to and activate the ATPase torsinA in the perinuclear space (33). Mice with hepatocyte-specific depletion of torsinA have even more profoundly decreased VLDL secretion and more severe steatosis than those with LAP1-depletion, but do not have nuclear LDs (32). Deficiency of LAP1, therefore, appears to have a unique role in nuclear LD generation. In the present study, we further characterize the nuclear LDs and investigate their mechanisms of formation in mouse hepatocytes with depletion of LAP1.

## MATERIALS AND METHODS

### Mice

The Columbia University Institutional Animal Care and Use Committee approved the protocols and procedures. The generation of mice with floxed alleles of the gene encoding Lap1 (*Tor1aip*^fl/fl^ or *Lap1*^fl/fl^) and breeding strategy for *Alb-Cre;Lap1*^fl/fl^ (L-CKO) mice with hepatocyte-specific depletion of LAP1 have been previously described (34). *Lmna*^fl/fl^ mice (35) were purchased from Jackson Laboratory (Stock number: 026284). The genetic background of all mice was C57BL/6J. Mice were housed in a climate-controlled room with a 12-hour light/12-hour dark cycle and fed regular chow diet (Purina Mills 5053) except when used in experiments testing the effects of a high fat diet. In those experiments, the high fat diet (Envigo, TD.88137) consisted of 42% of energy from fat (29% from saturated fat) and 30.5% from sucrose. We used both male and female mice for experiments on LAP1, as our previous studies showed no sexual dimorphism in nuclear LD formation with its depletion from hepatocytes (32). We also observed a similar frequency of nuclear LD-containing hepatocytes isolated from L-CKO mice from 2 months to less than 6 months of age; therefore, we used mice in this age range. We used only male mice for experiments examining depletion of lamin A/C, as mice with conditional depletion of these proteins from hepatocytes reportedly have sex-differences in liver fat accumulation (36).

### Primary hepatocyte isolation

Primary hepatocytes were isolated according to previously described methods (37). Briefly, livers were perfused *in situ* with Hank’s balanced salt solution without calcium containing 8 mM HEPES (Thermo Fisher Scientific) via the inferior *vena cava* after cutting the portal vein to allow outflow of the perfusate. They were perfused for 8 min at a rate of 5 ml/min at 37°C. This was followed by a perfusion at the same rate of Gibco DMEM, high glucose (Thermo Fisher Scientific, #11965118) with 80 mg/100 ml of collagenase type I (Worthington Biochemical Corporation) for 6 min. The livers were removed and minced in a Petri dish containing 4 ml of the same warm DMEM collagenase mixture for an additional 2 to 4 min. Ice-cold DMEM was added and the digested tissue filtered through nylon mesh and collected in a 50 ml conical tube. The suspension was centrifuged for 5 min at 50 x g. The supernatant was aspirated and the cell pellet washed 3 times with 30 ml of ice-cold DMEM. Viable cells were counted after staining with trypan blue. The isolated hepatocytes were plated onto collagen-coated 6-well plates at a density of 200,000 cells/well in 4 ml of DMEM plus 10% FBS and cultured for at least 2 hours prior to use for subsequent experiments.

### Cell culture and microscopy

Isolated hepatocytes were seeded onto collagen-treated coverslips in 6-well plates (150,000 cells/well) and grown over night at 37°C and 5% CO_2_ in high glucose DMEM (Thermo Fisher Scientific) containing 10% FBS (Thermo Fisher Scientific), 1% penicillin-streptomycin (Thermo Fisher Scientific) and 10 mM HEPES. FL83B cells (ATCC, CRL-2390) were cultured in Ham’s F-12K medium (Thermo Fisher Scientific #21127022) with 10% FBS. In some experiments, cells were incubated for 18-22 hours with 0.4 mM OA (Sigma-Aldrich) in complex with fatty-acid free BSA (Sigma-Aldrich) at a molar ratio of 6:1 added 4 hours after seeding. Primary Abs used were rabbit anti-adipose differentiation-related protein (ADRP) (Proteintech, 15294-1-AP) at a dilution of 1:100, rabbit anti-CCTα (Abcam, ab109263) at a dilution of 1:50, rabbit anti-lamin A/C (Abcam, ab133256) at a dilution of 1:200, rabbit anti-lamin B1 (38) at a dilution of 1:1,000, mouse anti-nuclear pore complex mAb414 (Abcam, ab24609) at a dilution of 1:100, rabbit anti-Sun2 (Abcam, ab87036) at a dilution of 1:500, rabbit anti-translocon-associated protein α (TRAPα) (39) at a dilution of 1:50, mouse anti-LBR (Abcam, ab232731) at a dilution of 1:250, and rabbit anti-PML protein (Abcam, ab53773) at a dilution of 1:100. FITC-conjugated goat anti-mouse and rhodamine-conjugated goat anti-rabbit secondary Abs were from Jackson ImmunoResearch Laboratories. Confocal fluorescence microscopy, imaging analysis, orthogonal and 3D reconstructions were performed on a Nikon A1 RMP microscope using NIS-elements software (Nikon) at the Confocal and Specialized Microscopy Shared Resource of the Herbert Irving Comprehensive Cancer Center at Columbia University Irving Medical Center. Widefield fluorescence microscopy was performed using a 40X Plan Apo objective (NA 1.0) on a Nikon Eclipse Ti microscope controlled by NIS-elements software. Acquired fluorescent images were processed using NIS-elements software, Image J or Fiji (40, 41).

### Transmission electron microscopy

Mouse livers were excised, cut into small pieces (1-2 mm), drop-fixed with 1% paraformaldehyde /2.5% glutaraldehyde in 0.1 M cacodylate buffer (pH 7.4) (Electron Microscopy Sciences), postfixed with 1% osmium tetroxide in 0.1 M cacodylate buffer (pH 7.4) (Electron Microscopy Sciences), incubated with uranyl acetate (Electron Microscopy Sciences) and dehydrated with ethanol. Tissues were subsequently rinsed with propylene oxide and embedded. Sections 60 nm thick were counterstained with uranyl acetate and lead citrate and examined using a JEM-1200EX electron microscope (JEOL).

### Quantification of nuclear LDs

Hepatocytes were plated onto 3 coverslips and lipid droplets stained with BOPIDY 493/503 (Thermo Fisher Scientific), LipiDye (Funakoshi) or Lipitox (Thermo Fisher Scientific) according to the manufacturers’ instructions. DAPI (Thermo Fisher Scientific) was used for nuclear staining. Micrographs of at least 3-5 different regions were taken from each stained coverslip. For quantification of nuclear LDs from images obtained by widefield microscopy, Image J with Cell Counter plugin was used to count the percentage of hepatocyte nuclei containing nuclear LDs. For a given image, 4 different groups were recorded: 1. nuclei containing 0-1 nuclear LDs, 2. nuclei containing 2-5 nuclear LDs, 3. nuclei containing 6-10 nuclear LDs, 4. nuclei with more than 10 nuclear LDs. Means and SEMs were calculated from the micrographs of each coverslips, then transferred to Microsoft Excel or GraphPad Prism (Ver 7) for graph generation and statistical analysis. For quantification from images obtained by confocal microscopy, nuclear LDs were counted manually from single-slice confocal images and analyzed using GraphPad Prism (Ver 7). We excluded small populations of cells negative for any lipid labeling, as those could have been non-hepatocytes. Hepatocytes were also stained with DAPI and micrographs analyzed using Image J to calculate nuclear Feret diameter and area.

### Adeno-associated virus (AAV)-mediated depletion of LAP1 and lamin A/C from hepatocytes

AAV8 vectors containing a control expression cassette (pAAV.TBG.PI.LacZ.bGH, plasmid 105534 [AAV-LacZ]) or encoding Cre recombinase (pAAV-TBG.PI.Cre.rBG, plasmid 107787 [AAV-Cre]) were gifts from James M. Wilson (University of Pennsylvania). Viral preparations were purchased from Addgene. *Lap1*^fl/fl^ and *Lmna*^fl/fl^ mice were randomly assigned to receive 1×10^11^ genome copies of AAV-LacZ or AAV-Cre via retro-orbital vein injection. Protein depletion was confirmed by immunoblotting with specific Abs against lamin A/C (Santa Cruz Biotechnology, sc-20681) at a dilution of 1:3000, LAP1 (33) at a dilution of 1:5000 and 7-actin (Santa Cruz Biotechnology, sc-47778) at a dilution of 1:3,000.

### Nuclear fractionation and lipidomic analysis

Primary hepatocytes isolated as described above were resuspended in hypotonic lysis buffer (10 mM HEPES, 5 mM MgCl_2_, 10 mM NaCl, pH 7.4) containing proteinase inhibitor cocktail (Sigma-Aldrich) and incubated on a rotating wheel for 30 minutes at 4°C. To isolate nuclei, the cell lysates were homogenized with a Wheaton Dounce homogenizer (DWK Life Sciences), underlaid with a 40% sucrose cushion, and centrifuged at 800 x g for 20 minutes. Fractionation was verified by immunoblotting using Abs against the nuclear proteins lamin A/C (Santa Cruz Biotechnology, sc-20681) at a dilution of 1:1,000 and the cytoplasmic protein GAPDH (Thermo Fisher Scientific, AM4300) at a dilution of 1:5,000). Lipidomic analysis of purified nuclei was performed at the Columbia University Biomarker Core Laboratory using an LC-MS/MS platform comprised of a 6490 Triple Quadrupole mass spectrometer integrated with a 1260 Infinity liquid chromatography system (Agilent Technologies).

### Antisense oligonucleotide (ASO) treatment

Mice were injected intraperitoneally twice a week with 50 mg/kg of an ASO against microsomal triglyceride transfer protein (MTP) (ISIS 144477) or a scrambled ASO (ISIS 299705) for 6 weeks. The ASOs were 20-mer phosphorothioate oligonucleotides containing 2′-O-methoxyethyl groups at positions 1-5 and 15-20 and were provided by Ionis Pharmaceuticals. After 6 weeks of administration, mice were sacrificed for hepatocyte isolation or liver tissue collection. Depletion of MTP was confirmed by immunoblotting with Abs against MTP (BD Biosciences, 612022) at a dilution of 1:2,000.

### Statistics

Unpaired 2-tailed Student’s *t* test were used to compare differences between 2 groups. Comparisons between 3 or more groups were performed using one-way ANOVA with Tukey’s multiple comparison test. Methods used for each experiment are provided in the figure legends. Statistical significance was set at *P* < 0.05.

## RESULTS

### Quantification and characterization of nuclear LDs in hepatocytes with LAP1 depletion

We previously reported that L-CKO mice with depletion of LAP1 from hepatocytes have reduced hepatic VLDL secretion, steatosis and nuclear LDs (32). To further characterize intranuclear LDs in hepatocytes isolated from these and control (*Lap1*^fl/fl^) mice, we stained them with BODIPY 493/503 to label neutral lipids and DAPI to identify nuclei. Widefield fluorescence microscopy showed that a small percentage of hepatocytes from control mice contained 1 or 2 nuclear LDs, which were similar to those described in previous studies of rodent hepatocytes (6, 9); however, hepatocytes from L-CKO mice contained frequent nuclear LDs, which were readily detectable by widefield fluorescence microscopy (Fig. 1A). Zoomed-in views showed that while some control hepatocytes had a few very small nuclear LDs, those from L-CKO mice contained variable numbers of different sizes, sometimes occupying nearly the entire nucleoplasmic area (Fig. 1B). While fewer than 4% of hepatocyte nuclei from control mice contained 2 or more nuclear LDs, nearly 60% of nuclei in hepatocytes from L-CKO mice did, with approximately 20% containing more than 10 (Fig. 1C). The nuclei in cultured hepatocytes from L-CKO mice had significantly increased Feret diameter and area (Fig. 1D). We corroborated the findings of nuclear LDs within larger nuclei of hepatocytes isolated from L-CKO mice by performing transmission electron microscopy on liver sections. LDs were inside larger nuclei in liver sections from L-CKO but not control mice (Fig. 1E). This was consistent with our previously published findings (32). These results indicated that hepatocytes isolated from L-CKO mice had larger nuclei containing nuclear LDs similar to intact liver tissue.

**Fig. 1.**
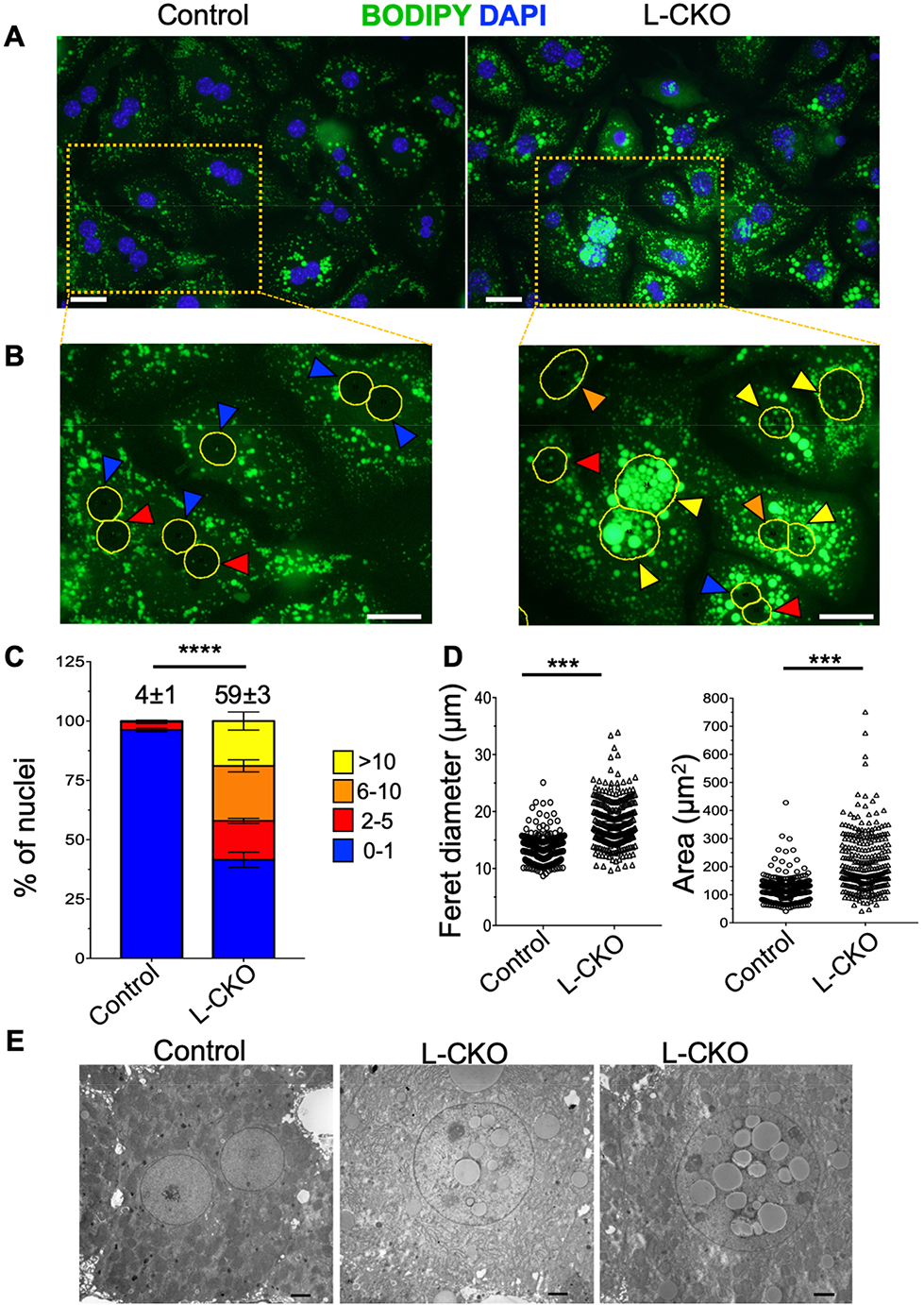
Nuclear LDs in hepatocytes isolated from L-CKO mice. (A) Representative widefield fluorescence photomicrographs of hepatocytes isolated from control (*Lap1*^fl/fl^) and L-CKO mice stained with BODIPY (green) and DAPI (blue). Scale bars: 25 μm. (B) Zoomed-in views of the areas within the orange rectangles in Panel A with nuclei outlined in yellow (based on DAPI labeling). Colored arrowheads indicate number of LDs per nucleus: blue 0-1 LDs, red 2-5, orange 6-10 and yellow >10. Scale bars: 25 µm. (C) Stacked column graphs with different colors representing the percentages of hepatocyte nuclei containing the indicated numbers of nuclear LDs. We analyzed a total of 604 (control) and 488 (L-CKO) nuclei of hepatocytes cultured on 3 different coverslips (n = 3 per genotype). The numbers at the top of the graphs indicate the mean percentages of hepatocyte nuclei with 2 or more nuclear LDs. These values and those within graphs are means ± SEM. \*\*\*\**P* < 0.0001 for percentage of hepatocyte nuclei with 2 or more nuclear LDs by 2-tailed Student’s *t* test. (D) Hepatocytes were stained with DAPI and Feret diameter (left panel) and area (right panel) of nuclei measured using Image J. Each dot represents a nucleus. 300 (control) and 279 (L-CKO) nuclei were plotted; \*\*\**P* < 0.001 by 2-tailed Student’s t test. (E) Transmission electron micrographs of liver sections showing nuclei from one control and two L-CKO mice. Scale bars: 2 µm.

To enhance lipid uptake, we cultured primary hepatocytes for 18-22 hours in media with 0.4 mM OA. Confocal fluorescence microscopy of these cells stained with BODIPY and DAPI showed that OA treatment led to an increase in the number and size of cytoplasmic LDs in hepatocytes from both control and L-CKO mice, with obvious nuclear LDs detected on zoomed-in views of hepatocytes from L-CKO mice, both with and without the addition of OA (Fig. 2A). However, OA treatment did not significantly increase the number of nuclei with LDs in control or L-CKO hepatocytes (Fig. 2B). These findings in primary hepatocytes are in contrast to those in cell lines where the addition of OA to culture media increases nuclear LD formation (9, 10). In this analysis using confocal fluorescence microscopy, we detected nuclear LDs in approximately 35% of hepatocyte nuclei from L-CKO mice in the absence of OA, compared to approximately 60% when we used wide-field fluorescence microscopy (see Fig. 1C). This is likely due to the fact that confocal microscopy takes images of a single optical section, which avoids counting cytoplasmic LDs overlaying a nucleus as nuclear LDs. However, it is possible that single confocal optical sections may miss a significant number of nuclear LDs present in other focal planes of the nucleus. To exclude this possibility, we used maximum intensity projections after taking Z-stacked confocal images to capture all LDs that appeared to be in a given nucleus. Single confocal optical sections from the center of a nucleus captured the majority of nuclear LDs seen in a maximum intensity projection image of the same nucleus (Fig 2C). The apparent nuclear LDs not seen in single confocal sections but captured in the maximum intensity projections were mostly located outside the nucleus or on its surface (Fig. 2D). Lipid droplets outside of, but touching the surface of the nucleus, were readily apparent on zoomed-in high-magnification images of orthogonal projections (Fig. 2E). Hence, using either maximum intensity projections or wide-field microscopy would likely lead to overcounting of nuclear LDs, as the counts would include some outside of the nucleus. Whether we used wide-field or confocal microscopy to count nuclear LDs in a given experiment, we always used the same imaging method and included controls for comparison. This assured that differences between the experimental and control groups were not a result of variations in the methods used to count nuclear LDs.

**Fig. 2.**
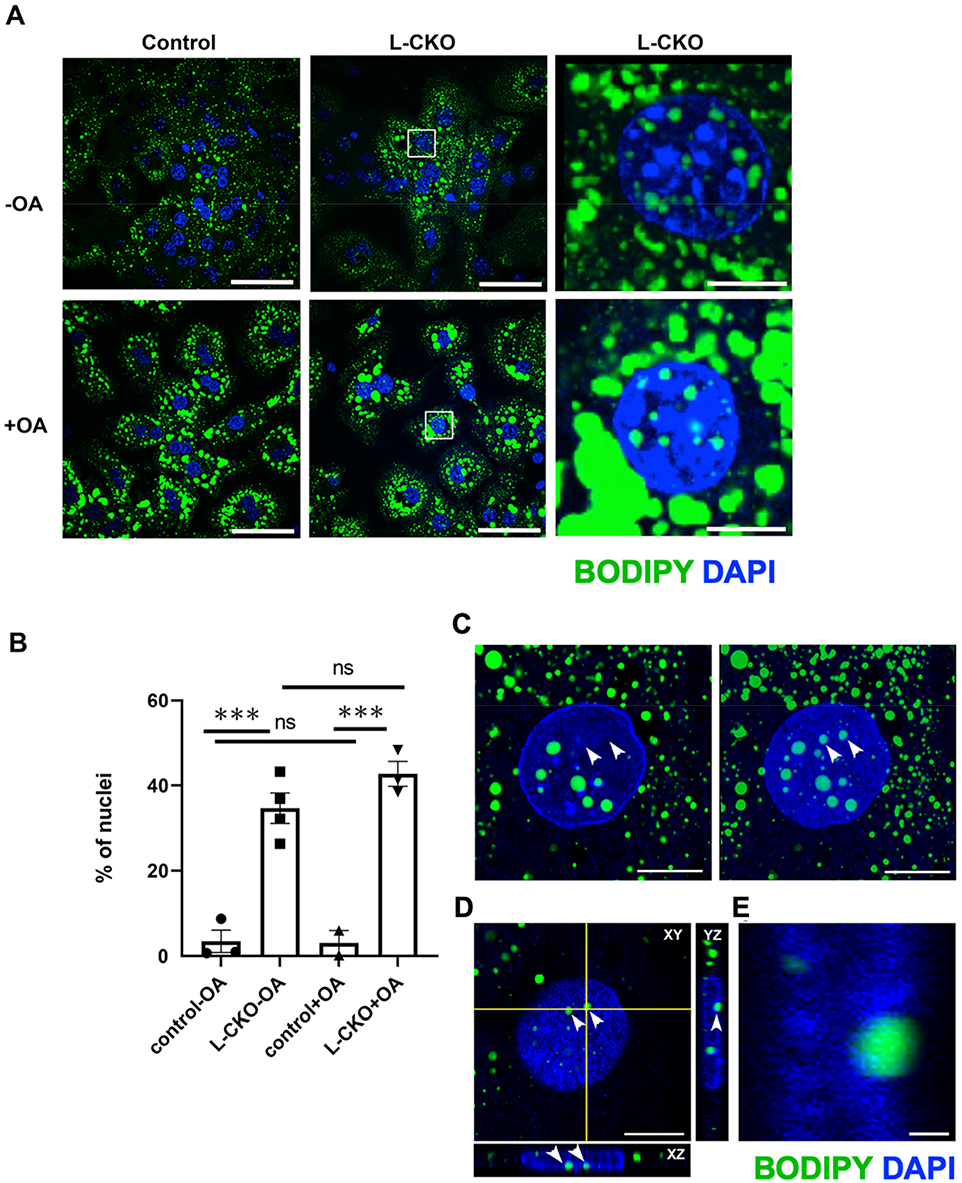
Effects of OA on LDs in hepatocytes from L-CKO mice. (A) Representative confocal fluorescence photomicrographs of hepatocytes from control (*Lap1*^fl/fl^) and L-CKO mice stained with BODIPY (green) and DAPI (blue). Panels to the right show zoomed-in views of the areas marked with white squares in the middle panels. Scale bars: 40 µm (left and middle panels) and 10 µm (right panels). (B) Percentages of hepatocytes from control and L-CKO mice, cultured without (-OA) or with (+OA) OA in the media, with nuclear LDs. Columns show mean percentage of nuclei with LDs, black symbols indicate data from separate experiments (> 170 nuclei counted for each experiment and condition) and error bars show SEM; ns = not significant, \*\*\**P* < 0.001 by one-way ANOVA with Tukey’s multiple comparison test. (C) A confocal single slice image from the center of a nucleus (left panel) captures the majority of nuclear LDs seen in a maximum intensity projection image of the same nucleus (right panel). Nuclear LDs not seen in the single slice photos but captured in the maximum intensity projection are indicated by arrowheads. Scale bar: 10 µm. (D) XY surface view of the nucleus shown in C and XZ and YZ orthogonal projections. Scale bar: 10 µm. (E) A zoomed-in view of the nuclear LD labeled with an arrow in the YZ projection in D, showing its location on the nuclear surface. Micrographs in panels C-E show hepatocytes from L-CKO mice labeled with BODIPY (green) and DAPI (blue). Scale bar: 1 µm.

Some proteins involved in lipid metabolism are associated with nuclear LDs in cultured cell lines, whereas other proteins that function in the formation of cytoplasmic LDs are not. CCT⍺ associates with nuclear LDs in U2OS and hepatocyte-derived cell lines, but ADRP (also known are perilipin-2), a component of cytoplasmic LDs, does not (9). We examined the colocalization of these proteins with nuclear LDs in hepatocytes from L-CKO mice. Confocal immunofluorescence microscopy revealed a diffuse nucleoplasmic localization of CCT⍺ in hepatocytes from control mice, whereas in some hepatocytes from L-CKO mice, especially after addition of OA to the media, there was no to minimal nucleoplasmic and some cytoplasmic localization (Fig. 3A). CCT⍺ was nucleoplasmic in nearly all control hepatocytes compared to about 80% of L-CKO hepatocytes without the addition of OA and only in approximately 40% of nuclei after OA addition (Fig. 3B). As nucleoplasmic CCT⍺ is reportedly inactive (14), the enzyme may therefore be active in the subset of hepatocytes from L-CKO that do not have it diffusely spread in the nucleus. After addition of OA to the culture media, 3D reconstructions from confocal images of hepatocytes from L-CKO mice showed that CCT⍺ was often concentrated around nuclear LDs and at the nuclear envelope, but was not robustly associated with cytosolic LDs (Fig. 3C). In contrast, ADRP was rarely colocalized with nuclear LDs but highly colocalized with cytosolic ones (Fig. 3D). These results show that nuclear and cytoplasmic LDs in hepatocytes of L-CKO mice differ in their associated proteins, similar to what has been shown in cultured cell lines (9).

**Fig. 3.**
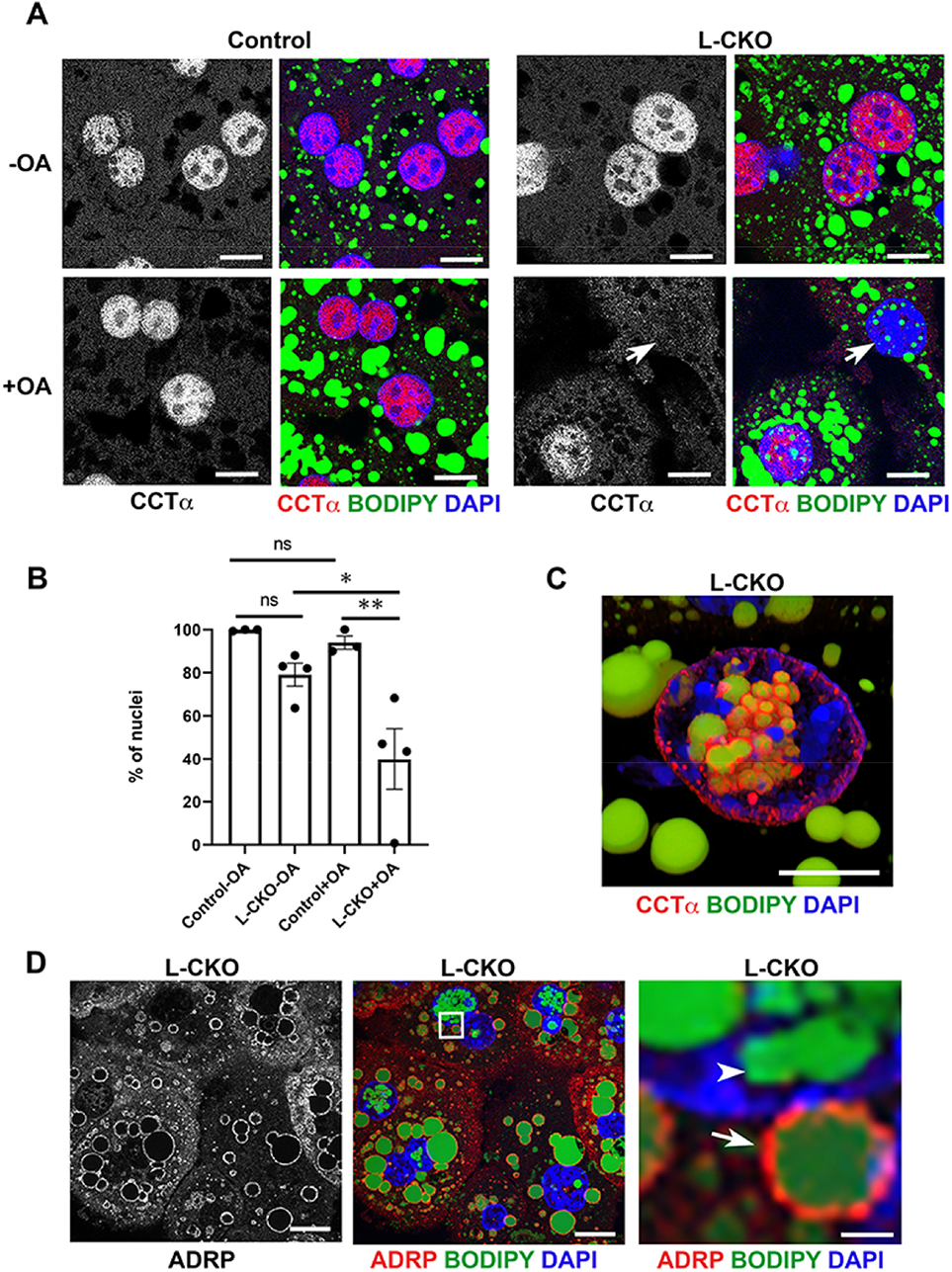
CCT⍺ and ADRP localization in hepatocytes from L-CKO mice. (A) Representative confocal photomicrographs of hepatocytes from control (*Lap1*^fl/fl^) and L-CKO mice, cultured with or without OA in the medium. The left panels from each genotype show labeling by anti-CCT⍺ Abs and the right panels show an overlay of labeling with anti-CCT⍺ Abs (red), BODIPY (green) and DAPI (blue). Arrows indicate a nucleus lacking nucleoplasmic CCTα. Scale bars: 10 µm. (B) Percentages of nuclei from control and L-CKO hepatocytes, cultured without (-OA) or with (+OA) OA in the media, with diffuse nucleoplasmic CCTα labeling detected by fluorescence microscopy. Columns show mean percentages of nuclei with diffuse nucleoplasmic CCTα localization, black circles indicate data from separate experiments (40-348 nuclei counted for each experiment and condition) and error bars show SEM. ns = not significant, \**P* < 0.05, \*\**P* < 0.01 by one-way ANOVA with Tukey’s multiple comparison test. (C) Representative 3D image reconstruction from confocal micrographs of a hepatocyte from an L-CKO mouse, cultured with OA in the media and labeled with anti-CCTα Abs (red), BODIPY (green), and DAPI (blue). Scale bar: 10 µm. (D) Representative confocal photomicrographs of hepatocytes from an L-CKO mouse; the left panel shows labeling with anti-ADRP Abs whereas the middle panel shows an overlay of labeling with anti-ADRP Abs (red), BODIPY (green) and DAPI (blue). The right panel shows a zoomed-in view of the area marked with a white square in the middle panel. Arrowhead indicates a nuclear LD lacking ADRP; arrow indicates a cytoplasmic LD with ADRP. Scale bars: 10 µm (left and middle panels) and 1 µm (right panel).

Nuclear LDs could form when phospholipid metabolism is disrupted. For example, excess cellular levels of phosphatidic acid lead to the formation of “supersized” LDs in yeast (43). We therefore asked if the levels of major phospholipid species are altered in the nuclei of hepatocytes from L-CKO mice. We isolated nuclei of hepatocytes from control and L-CKO mice by subcellular fractionation and validated the fractionation by immunoblotting with Abs against lamin A/C, a nuclear protein, and GAPDH, a cytosolic protein (Fig. S1A). We then measured phospholipid species in isolated nuclei by LC-MS/MS. There were no significant differences in the major phospholipid species measured, including phosphatic acid and phosphatidylcholine, or in the ratio of phosphatic acid to phosphatidylcholine (Table 1). However, the phosphatidylcholine to phosphatidylethanolamine ratio was significantly increased in nuclei of hepatocytes from L-CKO mice (Fig. S1B). There was also a significant increase of approximately 8-fold in triacylglycerol in nuclei of L-CKO mice to controls (Fig. S1C). This large increase in nuclear triacylglycerol assessed by chemical methods in isolated nuclei validated our microscopic analysis of BODIPY-stained cells showing increased intranuclear lipids in hepatocytes from L-CKO mice.

**Table 1.** Results from lipidomic analysis of nuclear fraction from hepatocytes from control and L-CKO mice (n=3 per group). The abbreviated and full names of lipid species are shown in the left two columns. The levels of each lipid species are represented as mol % of total lipid composition. The results from three different control and L-CKO mice are provided as well as means ± SEM (n = 3 per group). \*\*\**P* < 0.001, by unpaired Student’s *t* test.

| Abbreviation | Full name | Mol % of total lipid | | | Mean ± SEM | Mol % of total lipid | | | Mean ± SEM | P value |
| --- | --- | --- | --- | --- | --- | --- | --- | --- | --- | --- |
|  |  | control 1 | control 2 | control 3 |  | L-CKO 1 | L-CKO 2 | L-CKO 3 |  |  |
| SM | Sphingomyelin | 1.9440 | 1.2471 | 1.5860 | 1.5924 ± 0.201 | 1.7926 | 1.8697 | 3.0520 | 2.2381 ± 0.4075 | 0.2284 |
| PA | Phosphatidic acid | 0.2915 | 0.3262 | 0.2270 | 0.2815 ± 0.029 | 0.2615 | 0.3386 | 0.3846 | 0.3282 ± 0.0359 | 0.3697 |
| PC | Phosphatidylcholine | 3.8017 | 2.2881 | 2.5536 | 2.8811 ± 0.4666 | 3.5386 | 4.5045 | 5.6653 | 4.5694 ± 0.6148 | 0.094 |
| PE | Phosphatidylethanolamine | 19.3668 | 12.1711 | 13.1774 | 14.9050 ± 2.2505 | 11.1433 | 14.6894 | 16.9155 | 14.2493 ± 1.681 | 0.8268 |
| PS | Phosphatidylserine | 1.2359 | 0.7908 | 0.4832 | 0.8366 ± 0.2185 | 1.0402 | 0.9940 | 2.1940 | 1.4093 ± 0.3925 | 0.2713 |
| PI | Phosphatidylinositol | 1.8524 | 1.0944 | 1.3883 | 1.4450 ± 0.0202 | 1.3933 | 1.2974 | 2.3959 | 1.6951 ± 0.0022 | 0.5785 |
| TG | Triacylglycerol | 2.6072 | 2.4990 | 3.9703 | 3.0255 ± 0.4734 | 26.6863 | 19.7849 | 24.9933 | 23.8215 ± 2.077 | 0.0006*** |
| PC/PE | | 0.1963 | 0.1880 | 0.1938 | 0.1932 ± 0.0024 | 0.3176 | 0.3066 | 0.3349 | 0.3197 ± 0.0082 | 0.0001*** |
| PA/PC | | 0.0767 | 0.1426 | 0.0889 | 0.1027 ± 0.0036 | 10.7123 | 14.7780 | 7.7100 | 11.0667 ± 0.0021 | 0.4014 |

In cultured transformed hepatocyte cell lines, nuclear LDs are associated with PML nuclear bodies (9, 10). However, using Abs against PML protein for immunofluorescence microscopy, we could not detect PML nuclear bodies in hepatocytes from L-CKO mice, including those containing nuclear LDs, whereas we did detect them in FL83B cells, a transformed cell line derived from mouse fetal hepatocytes (42) (Fig. S2). Hence, unlike cultured hepatoma and osteosarcoma cell lines (9, 10), nuclear LDs in differentiated hepatocytes isolated from L-CKO are not associated with PML nuclear bodies.

### Nuclear LDs in hepatocytes with LAP1 depletion likely form from inner nuclear membrane invaginations

To determine how nuclear LDs form in hepatocytes from L-CKO mice, we examined their association with the nuclear envelope. Confocal microscopy of L-CKO hepatocytes double labeled with Abs against lamin A/C, an extrinsic protein of the inner nuclear membrane, and a lipid dye revealed nuclear envelope invaginations that colocalized with intranuclear lipid (Fig. 4A). We observed nuclear envelope invaginations in approximately 25% of hepatocyte nuclei from L-CKO mice (Fig. 4B). Nuclear envelope invaginations were also detected in hepatocytes from L-CKO mice labeled with Abs against lamin B1, another extrinsic protein of the inner nuclear membrane (Fig. S3). High-magnification orthogonal confocal views showed that some nuclear LDs in hepatocytes from L-CKO mice were very closely associated with the nuclear envelope invaginations labeled with Abs against the intrinsic inner nuclear membrane protein Sun2 (Fig. 4C). 3D reconstructions generated from confocal microscopic images of hepatocytes from L-CKO mice labeled with a lipid dye and anti-Sun2 Abs similarly revealed a close association with the nuclear envelope (Fig. 4D). Not all nuclear LDs were surrounded by lamins and Sun2, suggesting that as they enlarge they break free from the initially surrounding nuclear envelope invaginations.

**Fig. 4.**
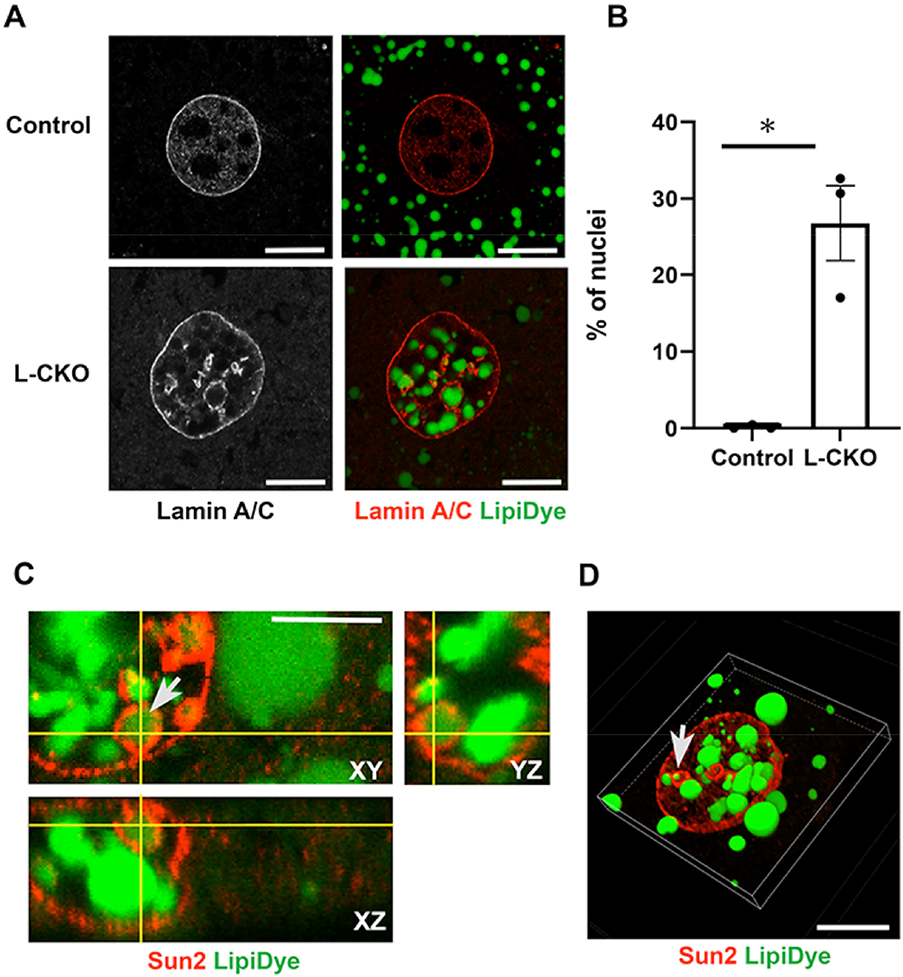
(A) Representative confocal photomicrographs in left panels show hepatocytes from control (*Lap1*^fl/fl^) and L-CKO mice labeled with anti-lamin A/C Abs and color confocal photomicrographs in right panels show an overlay of anti-lamin A/C Abs (red) and LipiDye (green) labeling. Note intranuclear lamin A/C and lipids in L-CKO hepatocytes. Scale bars: 10 µm. (B) Percentages of nuclei with nuclear envelope invaginations in hepatocytes from control and L-CKO mice. Columns show mean percentage of nuclei with nuclear invaginations, black circles indicate data from separate experiments (75-411 nuclei counted for each experiment and genotype) and error bars show SEM. \**P* < 0.05 by Student’s t test. (C) Representative orthogonal confocal sections showing XY, XZ and YZ planes of a partial view of a L-CKO mouse hepatocyte nucleus labeled with anti-Sun2 Abs (red) and LipiDye (green). Arrow indicates a LD inside a nuclear envelope invagination. Scale bar: 5 µm. (D) Representative 3D reconstruction of a hepatocyte nucleus from a L-CKO mouse, based on confocal microscopy data from labeling with anti-Sun2 Abs (red) and LipiDye (green). Arrow indicates a LD inside a nuclear envelope invagination. Scale bar: 10 µm.

We next determined if nuclear envelope invaginations in hepatocytes from L-CKO mice are type 1 or type 2 nucleoplasmic reticulum. Type 2 nucleoplasmic reticula are invaginations of both inner and outer nuclear membranes that contain nuclear pore complexes and type 1 are invaginations of only the inner nuclear membrane (17). Confocal fluorescence microscopy showed that nuclei of hepatocytes from L-CKO mice had considerable intranuclear labeling with anti-lamin A/C Abs; however, there was virtually no intranuclear labeling by an Ab that recognizes nuclear pore complex proteins (Fig. 5A). 3D image reconstructions clearly showed lamin A/C at the nuclear periphery as well as in an intranuclear reticular structure, whereas nuclear pore complexes were essentially only at the nuclear periphery (Fig. 5B). Confocal immunofluorescence microscopy of hepatocytes labeled with Abs against TRAP7, a protein of the ER/outer nuclear membrane, did not reveal nuclear invaginations (Fig. 5C). 3D image reconstruction of cells labeled with anti-TRAP1α Abs also did not reveal nuclear invaginations (Fig. 5D). These data indicate that nuclear LDs in hepatocytes depleted of LAP1 likely form from inner nuclear membrane invaginations (type 1 nucleoplasmic reticulum), which sometimes appear to surround nuclear LDs.

**Fig. 5.**
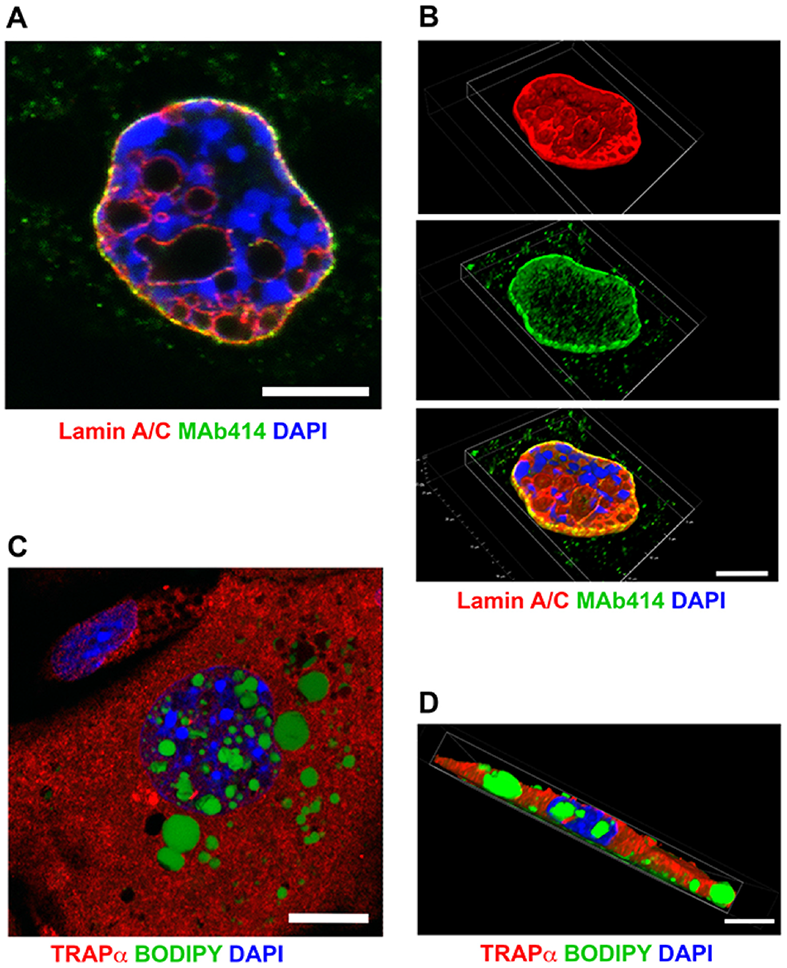
Nuclear envelope invaginations in hepatocytes from L-CKO mice are type 1 nucleoplasmic reticula. (A) Representative confocal fluorescence photomicrograph of a hepatocyte from a L-CKO mouse labeled with anti-lamin A/C Abs (red), Ab MAb414 that recognizes several nuclear pore complex proteins (green) and DAPI (blue); overlap of red and green appears yellow. Pore complexes are at the periphery and excluded from the interior of the nucleus. Scale bar: 10 µm. (B) 3D reconstruction of the nucleus shown in A labeled with anti-lamin A/C Abs (red), Ab MAb414 (green), and DAPI (blue). Top panel shows the lamin A/C signal (red), middle panel the pore complex signal (green) and bottom panel those signals plus DAPI (blue) together, with overlap of red and green appearing yellow. Lamin A/C is located in the nuclear interior and the periphery while pore complexes are only at the periphery and excluded from the interior of the nucleus. Scale bar: 10 µm. (C) Representative confocal fluorescence photomicrograph of a hepatocyte from a L-CKO mouse labeled with Abs against TRAPα, a marker of the ER/outer nuclear membrane (red), BODIPY (green) and DAPI (blue). TRAPα is essentially excluded from the nucleus. Scale bar: 10 µm. (D) Representative 3D reconstruction of a hepatocyte from a L-CKO mouse labeled with Abs against TRAPα, (red), BODIPY (green) and DAPI (blue). Scale bar: 10 µm.

### Differential effects of LAP1 and lamin A/C depletion on hepatocyte nuclear envelopes and nuclear LDs

LAP1 is associated with the nuclear lamina (44, 45). We therefore asked if a general structural perturbation of the nuclear envelope and lamina could cause nuclear LD accumulation in hepatocytes. A previous study reported that depletion of lamin A/C from mouse hepatocytes leads to male-specific hepatic steatosis (36). That study did not examine nuclear LDs. We therefore injected *Lmna*^fl/fl^ and *Lap1*^fl/fl^ male mice with AAV-Cre to induce depletion of the proteins from hepatocytes. *Lmna*^fl/fl^ mice were also injected with AAV-LacZ as a control. We confirmed efficient depletion of lamin A/C and LAP1 from hepatocytes of these mice 8 weeks after injection with AAV-Cre (Fig. 6A). Immunofluorescence microscopy of lamin A/C-depleted hepatocytes labeled with anti-lamin B1 Abs and DAPI revealed severely misshapen nuclei with deformed lamina-free parts of the nuclear envelope, which were not seen in hepatocytes with LAP1 depletion or from *Lmna*^fl/fl^ mice injected with AAV-LacZ (Fig. 6B). These misshapen nuclei were present in approximately 50% of hepatocytes with lamin A/C depletion, but only approximately 5% of those with LAP1 depletion (Fig. S4A). Labeling with Abs against LBR, an intrinsic protein of the inner nuclear membrane, showed that lamin A/C-depleted hepatocytes similarly had LBR-free areas of their nuclear envelopes, which were rarely present in hepatocytes from AAV-LacZ-injected *Lmna*^fl/fl^ mice (Fig. S4B). As with anti-lamin B1 Ab labeling, approximately 45% of hepatocytes lacking lamin A/C had misshapen nuclei when examined by anti-LBR Ab labeling (Fig. S4C). We stained lamin A/C-depleted hepatocytes with BODIPY and DAPI and performed fluorescence microscopy to identify nuclear LDs. Whereas many hepatocytes with LAP1 depletion had more than 2 nuclear LDs, sometimes many large ones, lamin A/C depletion led to only small LDs in fewer cells (Fig. 6C). Approximately 34% of lamin A/C-depleted hepatocyte nuclei had nuclear LDs whereas 69% of LAP1-depleted hepatocyte nuclei did (Fig. 6D). These results show that the depletion of lamin A/C from hepatocytes of adult mice causes severely misshapen nuclei and structural deformities of the nuclear envelope with areas devoid of key proteins. In these cells, small LDs may accumulate within the nuclei. In contrast, LAP1 depletion from hepatocytes does not cause severe nuclear envelope deformities but promotes more robust and frequent intranuclear LDs.

**Fig. 6.**
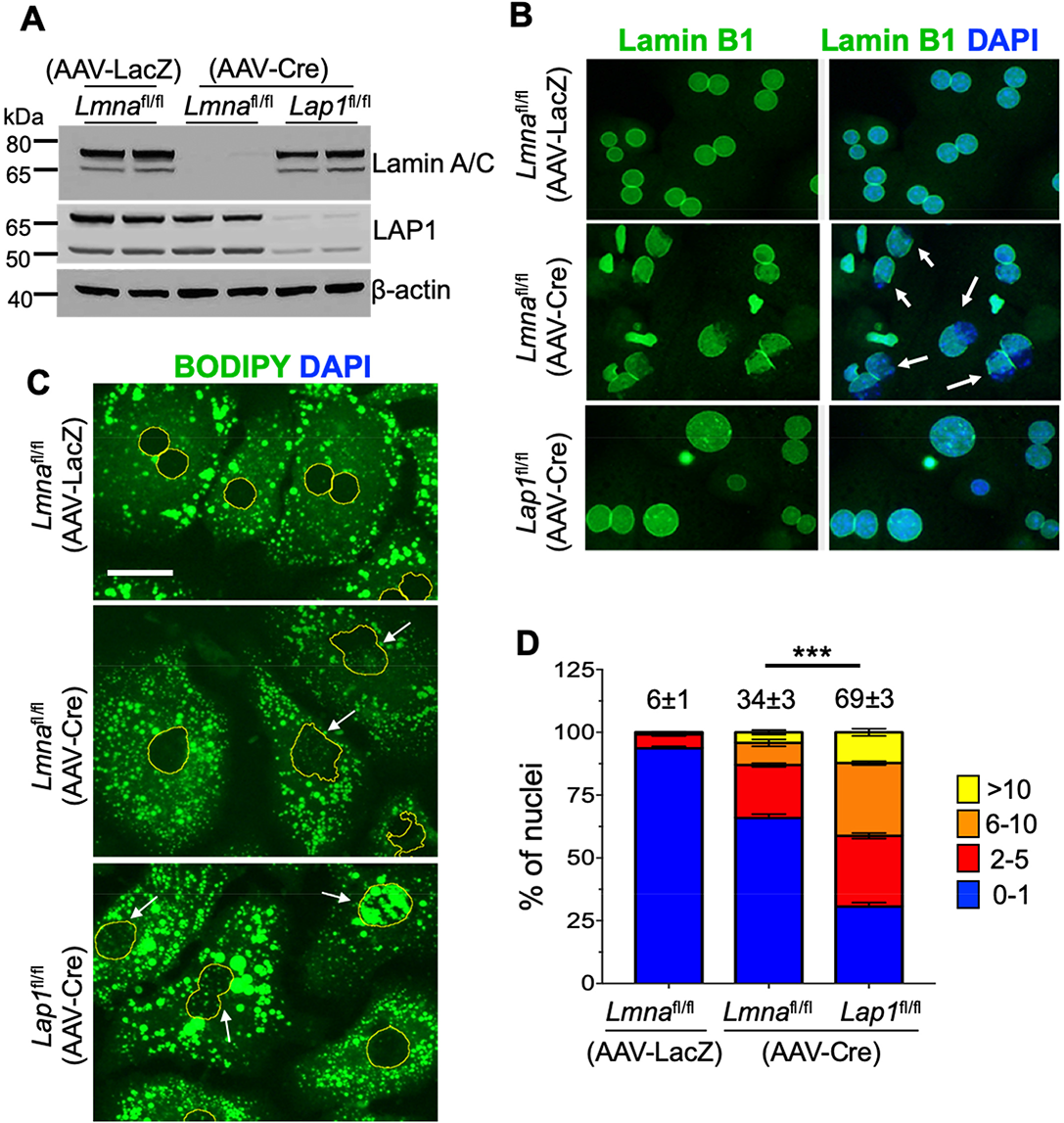
Hepatocytes isolated from male mice with acute depletion of lamin A/C or LAP1. (A) Immunoblots of hepatocyte protein lysates from male *Lmna*^fl/fl^ or *Lap1*^fl/fl^ mice 8 weeks after injection with AAV-LacZ or AAV-Cre, probed with Abs against lamin A/C, LAP1 and 7-actin. Each lane contains protein lysates from 2 different wells of hepatocytes isolated from one mouse for each condition. (B) Representative widefield immunofluorescence photomicrographs of hepatocytes stained with anti-lamin B1 Abs (green) in left panels and overlay with DAPI (blue) in right panels. Arrows indicate nuclei with absence of lamin B1 from parts of the nuclear envelope. Scale bar: 25 μm. (C) Representative widefield fluorescence photomicrographs of hepatocytes stained with BODIPY with nuclei outlined in yellow (based on DAPI labeling). Arrows indicate nuclei with nuclear LDs. Scale bar: 25 µm. (D) Stacked column graphs with different colors representing the percentages of hepatocyte nuclei containing the indicated numbers of nuclear LDs. We analyzed a total of 661 (AAV-LacZ injected *Lmna*^fl/fl^ mouse), 515 (AAV-Cre injected *Lmna*^fl/fl^ mouse) and 561 (AAV-Cre injected *Lap1*^fl/fl^ mouse) nuclei of hepatocytes cultured on 3 different coverslips (n=3 per group). The numbers at the top of the graphs indicate the mean percentages of hepatocyte nuclei with 2 or more nuclear LDs. These values and those within graphs are means ± SEM. \*\*\**P* < 0.001 for percentage of hepatocyte nuclei with 2 or more nuclear LDs by one-way ANOVA followed by Tukey’s multiple comparison test.

### Diet influences nuclear LDs in L-CKO mice

LDs are dynamic structures influenced by nutritional status (2, 46). Previous studies using hepatoma and U2OS cell lines demonstrated that culturing in media containing OA induced both cytosolic and nuclear LDs (9, 10), however we did not observe increased nuclear LDs upon OA treatment in hepatocytes isolated from L-CKO mice. We therefore asked if diet could modulate the frequency and size of nuclear LDs in hepatocytes from L-CKO mice. We initially hypothesized that feeding L-CKO mice a high fat diet would increase nuclear LD formation in hepatocytes and that fasting would lead to opposite results. L-CKO mice gained body mass at a similar rate as the control mice after 8 weeks on a high fat diet (Fig. 7A). However, the liver to body mass ratio was significantly greater in L-CKO compared to control mice (Fig. 7B). Histological examination of liver sections from high fat-fed L-CKO mice revealed increased macrovesicular steatosis (Fig. S5). Wide-field fluorescence microscopy of hepatocytes stained with BODIPY and DAPI showed an overall increase in cytoplasmic LDs in cells from L-CKO mice fed a high fat diet but fewer nuclear LDs (Fig. 7C). Refuting our initial hypothesis, only 14% of L-CKO mice fed a high fat diet had 2 or more nuclear LDs in their hepatocyte nuclei compared to 60% in mice fed a chow diet (Fig. 7D). In control hepatocytes, there was no significant difference in the number of nuclear LDs between hepatocytes from mice fed a chow or a high fat diet (Fig. S6). In a converse experiment, we examined the changes in nuclear LDs in hepatocytes isolated from chow-fed L-CKO mice after a 24-hour fast. Fasting promotes adipose tissue lipolysis with a subsequent increase in plasma free fatty acids, which in turn increases intrahepatic lipid content (47). Histological examination of liver sections showed slightly increased steatosis after 24 hours of fasting (Fig. S7) and hepatocytes isolated from L-CKO mice after fasting had more prominent cytosolic LDs but apparently fewer nuclear LDs than fed mice (Fig. 7E). Only 24% of fasted L-CKO mice had 2 or more nuclear LDs in their hepatocytes compared to 66% in mice fed a normal chow diet (Fig. 7F). In control mice, we observed no changes of the frequency of nuclear LDs after fasting (Fig. S8). In summary, both high fat diet and fasting, which cause an increase in overall cellular lipids by increasing delivery of exogenous or endogenous fat to the liver, respectively, reduced nuclear LDs in L-CKO mice.

**Fig. 7.**
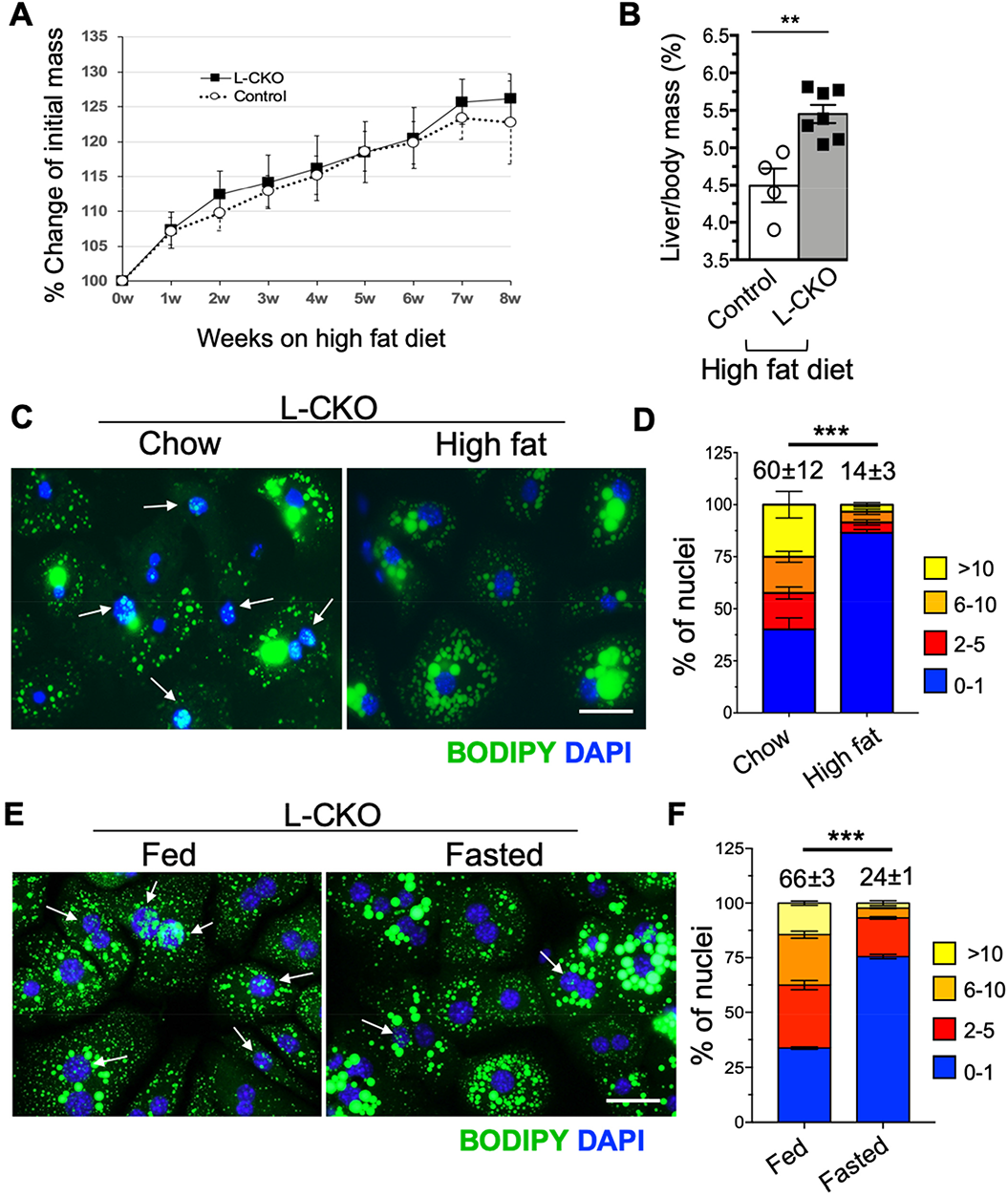
Analysis of nuclear LDs in hepatocytes isolated from high fat diet-fed and fasted L-CKO mice. (A) Percent change of initial body mass of mice while on a high fat diet. Control (*Lap1*^fl/fl^) and L-CKO mice at 8 weeks of age were fed a high fat diet for 8 weeks and body mass was measured weekly; n = 5-7 mice per group. Each circle and square indicate mean values of body mass. Error bars indicate SEM. (B) Liver to body mass ratios of control and L-CKO mice after 8 weeks on a high fat diet; n = 4-7 mice per group. Each circle and square indicate value from an individual mouse, rectangular bars show means and error bars indicate SEM; \*\**P* < 0.01 by 2-tailed Student’s *t* test. (C) Representative widefield fluorescence photomicrographs of hepatocytes from L-CKO mice stained with BODIPY (green) and DAPI (blue). Hepatocytes were isolated from chow-fed (left panel) or high fat diet-fed (right panel) L-CKO mice. Arrows indicate nuclei containing LDs. Scale bar: 25 μm. (D) Stacked column graph with different colors representing the percentages of hepatocyte nuclei containing the indicated numbers of nuclear LDs. We analyzed a total of 112 (chow diet) and 179 (high fat diet) nuclei of hepatocytes cultured on 3 different coverslips (n=3 per group). The numbers at the top of the graphs indicate the mean percentages of hepatocyte nuclei with 2 or more nuclear LDs. These values and those within graphs are means ± SEM. \*\*\**P* <0.001 for percentage of hepatocyte nuclei with 2 or more nuclear LDs by 2-tailed Student’s *t* test. (E) Representative widefield fluorescence photomicrographs of hepatocytes from L-CKO mice stained with BODIPY (green) and DAPI (blue). Hepatocytes were isolated from mice fed a normal chow diet (left panel) or mice fasted for 24 hours (right panel). Arrows indicate nuclei containing LDs. Scale bar: 25 μm. (F) Stacked column graph with different colors representing the percentage of nuclei containing the indicated numbers of nuclear LDs in hepatocytes isolated from L-CKO mice fed normally or after 24 hours of fasting. We analyzed a total of 479 (fed) and 599 (fasted) nuclei of hepatocytes cultured on three different coverslips (n=3 per group). The numbers on the top of the graphs indicate the mean percentages of hepatocyte nuclei with 2 or more nuclear LDs. Data are shown as means ± SEM. \*\*\**P* < 0.001 for percentage of hepatocyte nuclei with 2 or more nuclear LDs by 2-tailed Student’s *t* test.

### MTP depletion does not change the nuclear LD frequency in hepatocytes from L-CKO mice

An MTP inhibitor suppressed OA-induced nuclear LD accumulation in hepatocarcinoma cell lines (11). Therefore, we tested if the depletion of MTP could change the number of nuclear LDs in hepatocytes from L-CKO mice. To deplete MTP, we used a previously-described ASO (37). After 6 weeks of treatment, immunoblotting showed that this ASO led to efficient knockdown of MTP in hepatocytes from control and L-CKO mice, whereas a scrambled control ASO had no effect (Fig. 8A). Fluorescence microscopy of isolated hepatocytes stained with BODIPY and DAPI showed that MTP depletion increased the apparent number and size of cytosolic LDs in both control and L-CKO mice (Fig. 8B). However, MTP depletion did not change the percentages of nuclei with LDs (Fig.8C). These results indicate that the formation of nuclear LDs in L-CKO mice is independent of MTP activity.

**Fig. 8.**
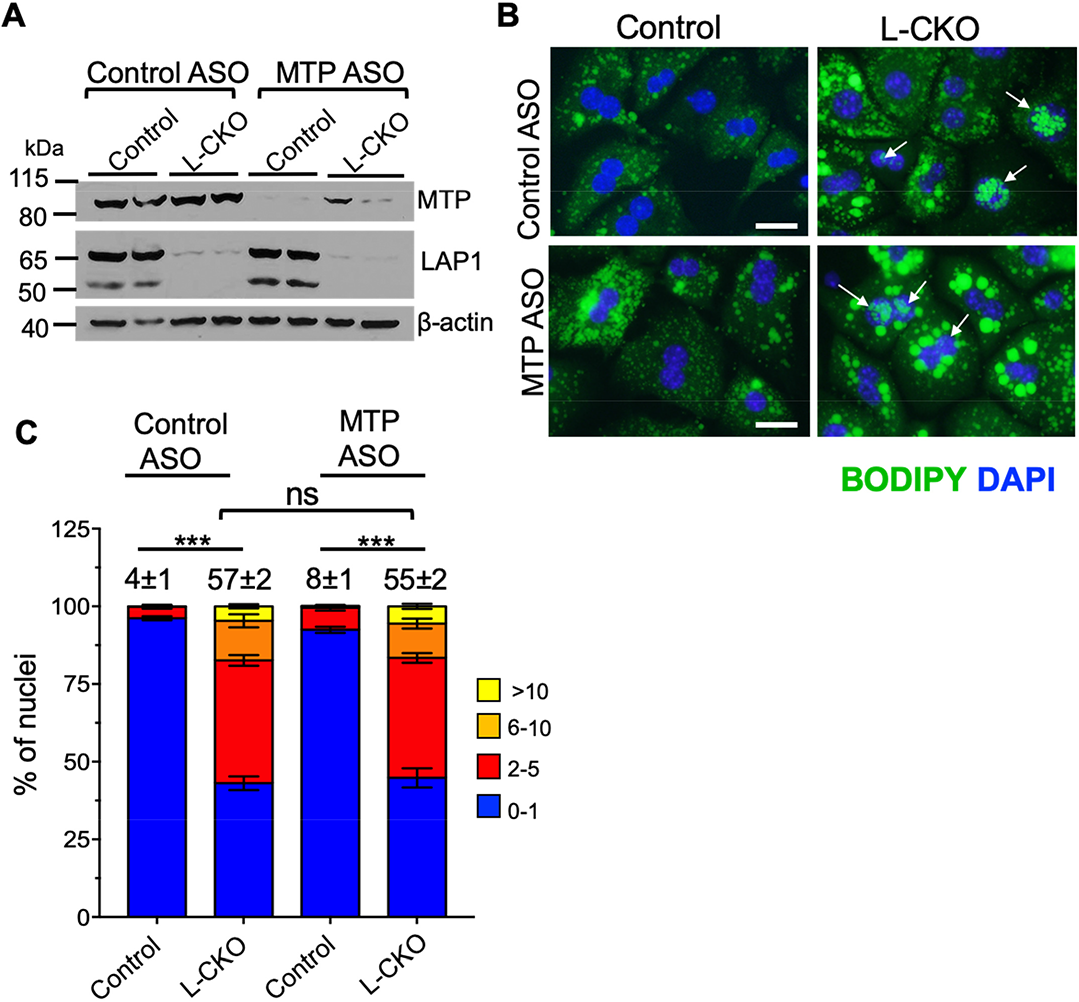
Effect of MTP depletion on nuclear LDs in hepatocytes from control and L-CKO mice. (A) Immunoblots of hepatocyte protein lysates from control and L-CKO mice after 6 weeks of scrambled control or MTP ASO treatment probed with Abs against MTP, LAP1 and 7-actin. Each lane contains protein lysates from 2 different wells of hepatocytes isolated from one mouse for each condition. (B) Representative widefield fluorescence photomicrographs of hepatocytes from control or L-CKO mice stained with BODIPY (green) and DAPI (blue). Hepatocytes were isolated from indicated mice after administration of control ASO (upper panels) or MTP ASO (lower panels). Arrows indicate nuclei containing LDs. Scale bars: 25 μm. (C) Stacked column graph with different colors representing the percentage of nuclei containing the indicated numbers of nuclear LDs. We analyzed a total of 774 (control mice given control ASO), 368 (L-CKO mice given control ASO), 511 (control mice given MTP ASO) and 462 (L-CKO mice given MTP ASO) from hepatocytes cultured on 3 different cover slips (n = 3 per group). The numbers at the top of the graphs indicate the mean percentages of hepatocyte nuclei with 2 or more nuclear LDs. These values and those within graphs are means ± SEM. ns = not significant by 2-tailed Student’s *t* test.

## DISCUSSION

Depletion of nuclear envelope protein LAP1 from hepatocytes *in vivo* leads to robust formation of nuclear LDs in chow-fed L-CKO mice. In hepatocytes isolated from these mice, the frequency of nuclei containing 2 or more LDs ranged from 30% to 70%, depending on the microscopic approach used to count them. In some LAP1-deficient hepatocytes, LDs occupied nearly the entire nuclear area. Only 4% of nuclei in hepatocytes isolated from control mice had LDs and they were much smaller than most of the LDs in LAP1-depleted hepatocytes.

In LAP1-deficient hepatocytes from L-CKO mice, nuclear LDs are often associated with nuclear lamins, whereas Ohsaki et al. (9) reported that lamins are depleted in cultured U2OS and hepatoma cells containing nuclear LDs. Nuclear LD formation is promoted in these cultured cell lines by adding OA to the medium (9, 10). However, overnight incubation of hepatocytes isolated from L-CKO mice in medium containing OA did not change the number of nuclear LDs, while cytosolic lipid content was increased. Despite these differences, nuclear LDs in LAP1-depleted hepatocytes and hepatoma cells are both associated with CCT⍺ and lack ADRP (9).

We demonstrated that LAP1-depleted hepatocytes have invaginations of the inner nuclear membrane, which have been termed type 1 nucleoplasmic reticulum (17). These invaginations are sometimes co-localized with nuclear LDs. Nuclear envelope invaginations also occur in fibroblasts from humans with a genetic mutation leading to complete deletion of LAP1 (48), suggesting that the protein plays an important role in maintaining nuclear integrity in cells other than hepatocytes. Nuclear LDs are also associated with type 1 nucleoplasmic reticula in hepatoma cell lines (9, 11). In these cultured cell lines, depletion of Sun proteins, which like LAP1 are monotopic integral inner nuclear membrane proteins, also leads to type 1 nucleoplasmic reticula and nuclear LD formation (9). However, we showed that lamin A/C-depletion from adult mouse hepatocytes *in vivo* resulted in disruption of nuclear envelope structure but few nuclear LDs. Hence, specific inner nuclear membrane invaginations, rather than a generalized disruption of the nuclear envelope, may play a critical role in nuclear LD formation.

In cultured hepatoma cell lines, nuclear LDs and type 1 nucleoplasmic reticula are associated with PML nuclear bodies (9, 10). In these cell lines, knockdown of PML isoform II decreases both nuclear LDs and type 1 nucleoplasmic reticula, whereas overexpression of the protein increases both (9). However, we could not detect PML nuclear bodies in differentiated hepatocytes isolated from L-CKO mice, including those containing nuclear LDs. In contrast, we could readily detect nuclear PML nuclear bodies in a transformed fetal mouse hepatocyte-derived cell line. This is consistent with the previously-reported absence or very-low-level expression of PML protein in normal hepatocytes that increases with inflammation, cirrhosis, dysplasia, and transformation to carcinoma (49–51). The minimal PML protein expression in L-CKO hepatocytes could not account for the formation of numerous large LDs in the nuclei of many cells. Therefore, unlike in transformed cultured cell lines, PML nuclear bodies do not appear to function in nuclear LD formation in differentiated hepatocytes depleted of LAP1. However, type 1 nucleoplasmic reticula appears to play a role in both.

Nuclear LDs in LAP1-deficient hepatocytes dynamically respond to the nutritional status of the mice from which they are isolated. A high fat diet and prolonged fasting both promote free fatty acid loading of hepatocytes, which can induce cytosolic LD formation (52). While cytosolic LDs were increased in hepatocytes of L-CKO mice after 8 weeks on either a high fat diet or 24 hours of fasting, nuclear LDs were significantly reduced. A recent study using lipid saturation sensors in *Saccharomyces cerevisiae* suggests that increased unsaturation of lipid enhances cytosolic LD formation while suppressing it in the nucleus (20). Future lipid profiling experiments of nuclear and cytoplasmic LDs in hepatocytes of L-CKO mice receiving different diets could provide insight regarding factors that influence their subcellular distribution in mammalian cells.

Our previous studies demonstrated that depletion of LAP1 from hepatocytes leads to reduced apoB100 and defective VLDL secretion (32). Soltysik et al. (11) proposed that nuclear LDs are derived from apoB-free VLDL precursors assembled in the ER lumen by MTP. In LAP1-depleted hepatocytes, increased expression or enhanced MTP activity with concurrent reduced availability of apoB100 could therefore be a possible mechanism of nuclear LD formation. However, when we depleted MTP from LAP1-deficient and control hepatocytes, we observed an apparent increase in the number and size of cytosolic LDs but no difference in percentages of hepatocytes with nuclear LDs. This shows that nuclear LDs in LAP1-deficient hepatocytes can get into the nucleus or be synthesized from the nuclear membrane independent of MTP. These results again further demonstrate that the nuclear LDs found in differentiated LAP1-deficient hepatocytes isolated from intact mouse livers differ from those observed in hepatocarcinoma lines.

We observed a reduced percentage of LAP1-depleted hepatocytes with nucleoplasmic, inactive CCT⍺. However, presumably-active CCT⍺ colocalized with nuclear LDs and the nuclear envelope, implying that LAP1 depletion leads to increased CDP-choline synthesis. There was also an increase in the phosphatidylcholine to phosphatidylethanolamine ratio in nuclei of hepatocytes from L-CKO mice compared to control mice, consistent with increased CDP-choline synthesis. Increased CDP-choline synthesis could also be a compensatory mechanism for reduced VLDL secretion, as these two processes are linked in cells (53–55). The potential role of increased CDP-choline synthesis, particularly at the nuclear envelope, therefore warrants future investigation of it as a potential factor in nuclear LD formation in LAP1-depleted hepatocytes.

In conclusion, our results demonstrate that LAP1-depleted hepatocytes from L-CKO mice provide a valuable tool to study the biogenesis and significance of nuclear LDs in mammalian liver. They also provide a mammalian model system to further study the subcellular partitioning of LDs in response to nutritional and metabolic alterations *in vivo*. In addition, as L-CKO mice develop steatosis and nonalcoholic steatohepatitis on a chow diet (32), they could also provide a valuable model to investigate the potential pathophysiological impact of nuclear lipid metabolism in these liver diseases.

## Data Availability

All data reported in this study are located within the main text or supplemental data and are available upon request from Dr. Ji-Yeon Shin (Columbia University).

## Abbreviations

AAV: adeno-associated virus
AAV-Cre: pAAV-TBG.PI.Cre.rBG
AAV-LacZ: pAAV.TBG.PI.LacZ.bGH
ADRP: adipose differentiation-related protein
ASO: antisense oligonucleotide
CCT⍺: CTP:phosphocholine cytidylyltransferase ⍺
LAP1: lamina-associated polypeptide 1
L-CKO mice: *Alb-Cre;Tor1aip1*^fl/fl^ mice
LDs: lipid droplets
MTP: microsomal triglyceride transfer protein
OA: oleic acid
PML: promyelocytic leukemia
TRAPα: translocon-associated protein α

## Acknowledgements

We thank Kristy Brown (Columbia University) for help with electron microscopy, and Renu Nandakumar and Yimeng Xu (Columbia University Biomarker Core Laboratory) for lipidomic analysis, Christopher Nicchitta (Duke University) for providing anti-TRAPα Abs and James Wilson (University of Pennsylvania) for AAV vectors. We also thank Weijia Fan (Columbia University Irving Institute for Clinical and Translational Research Biostatistics, Epidemiology and Research Design Core) for help with statistical analysis.

## Supplemental data

This article contains supplemental data.

## Author contributions

Conceptualization: H. N. G, H. J. W., J.-Y. S.; Methodology: C. Ö., A. H.-O., J.-Y .S.; Validation: C. Ö., A. H.-O., J.-Y. S.; Formal analysis: C. Ö., S. J. T., J.-Y. S.; Investigation: C. Ö., A. H.-O., S. J. T., J.-Y. S.; Resources: W. T. D., H. N. G., H. J. W., J.-Y. S.; Writing – original draft preparation: C. Ö., J. Y.-S.; Writing – review and editing: H. N. G., H. J. W.; Visualization: C. Ö., J.-Y. S.; Supervision: H. N. G., H. J. W., J.-Y. S; Project administration: H. J. W.; J.-Y. S; Funding acquisition: W. T. D., H. N. G., H. J. W.; J.-Y. S.

## Funding and additional information

This work was supported by a Columbia University Digestive and Liver Diseases Research Center Pilot Grant (J.-Y. S.); a Gilead Science Research Scholar Award (J.-Y. S.); an AASLD Pinnacle Research Award in Liver Disease (J.-Y. S); and the National Institutes of Health [grant numbers R01DK118480 (W. T. D., H. N. G., H. J. W.), R35HL135833 (H. N. G.)]. Some of the image processing and analysis was performed in the Confocal and Specialized Microscopy Shared Resource of the Herbert Irving Comprehensive Cancer Center at Columbia University, which is supported by the National Institutes of Health [grant number P30CA013696]. The Columbia University Irving Institute for Clinical and Translational Research Biostatistics, Epidemiology and Research Design Core, which provided help with statistics, is also supported by the National Institutes of Health [grant number UL1TR001873]. The content is solely the responsibility of the authors and does not necessarily represent the official views of the National Institutes of Health.

