## Supplemental materials for "Hepatocytes deficient in nuclear envelope protein lamina-associated polypeptide 1 are an ideal mammalian system to study intranuclear lipid droplets"

Supplemental Figure S1

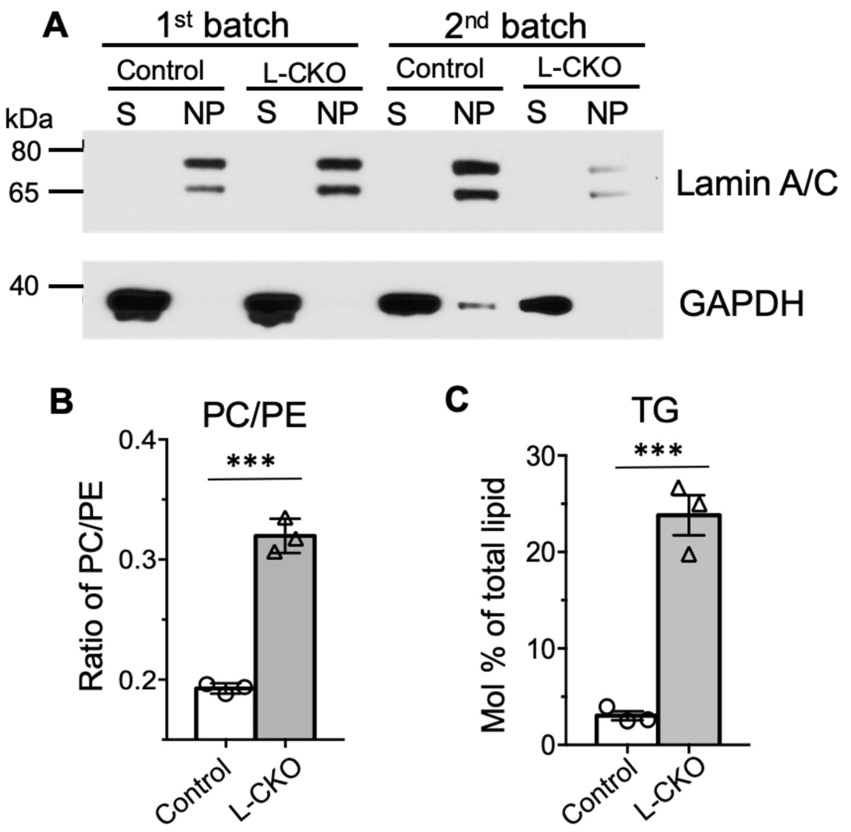


**Fig. S1.** Fractionation of hepatocyte nuclei and select results of lipidomic analysis. (A) Immunoblot showing cytosolic supernatants (S) and nuclear pellets (NP) from hepatocytes isolated from control and L-CKO mice on two separate dates (batch 1 and 2, respectively). Lamin A/C, a nuclear protein, is detectable in NP and GADPH, a cytosolic protein, in the S. (B) Phosphatidylcholine (PC)/phosphatidylethanolamine (PE) ratios of nuclear fractions from hepatocytes of control and L-CKO mice. Columns show mean PC/PE ratios, symbols indicate data from individual mice and error bars SEM (n = 3) ***$P < 0.001$ by 2-tailed Student's *t* test (C) Triacylglycerol (TG) content of nuclear fractions from hepatocytes of control and L-CKO mice. Columns show mean TG mol % of total lipid, symbols indicate data from individual mice and error bars SEM (n=3 per group). ***$P < 0.001$ by 2-tailed Student's *t* test.

Supplemental Figure S2

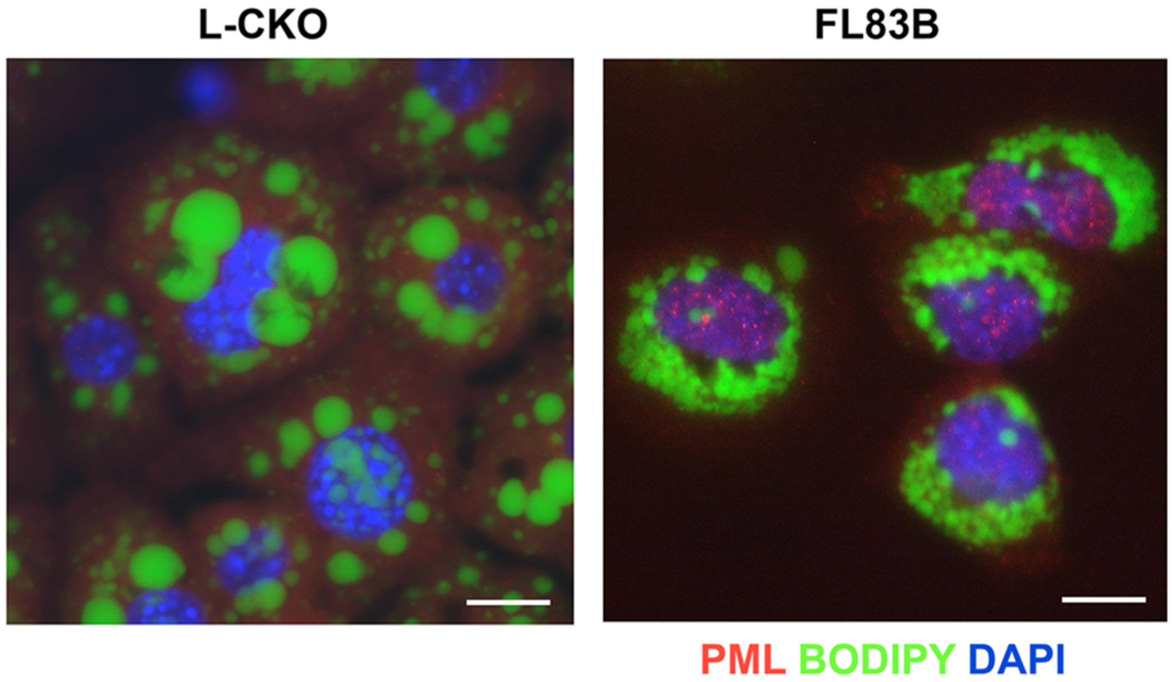


**Fig. S2.** Widefield fluorescence photomicrographs of hepatocytes from a L-CKO mouse (left panel) and FL83B cells (right panel), both cultured with OA in the media, labeled with anti-PML Abs (red), BODIPY (green) and DAPI (blue). PML labeling is detected as intranuclear dots in FL83B cells but not in L-CKO hepatocytes. Scale bars: 20 µm.

Supplemental Figure S3

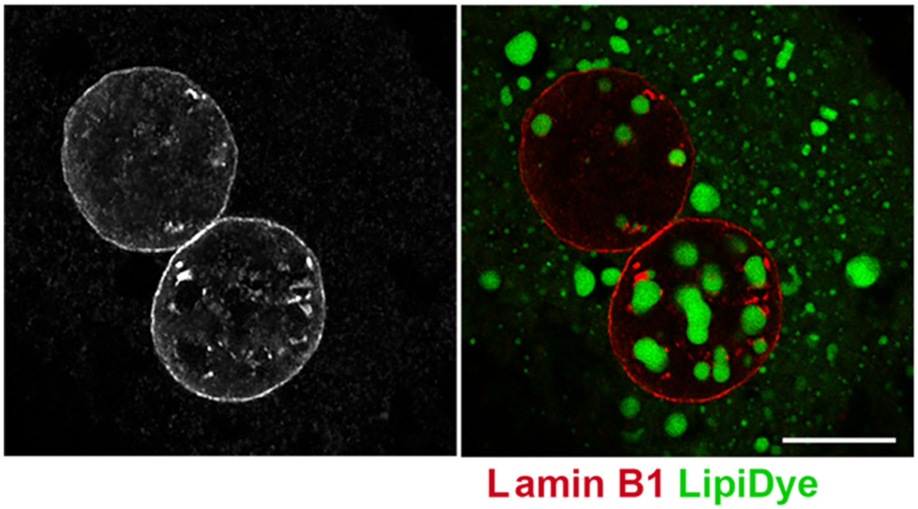


**Fig. S3.** Confocal photomicrographs of a hepatocyte from a L-CKO mouse labeled with anti-lamin B1 Abs and LipiDye showing nuclear envelope invaginations and nuclear LDs. Black and white micrograph in the left panel shows labeling with anti-lamin B1 Abs and color micrograph in the right panel an overlay of anti-lamin B1 Abs (red) and LipiDye (green) labeling. Scale bar: 10 μm.

Supplemental Figure S4

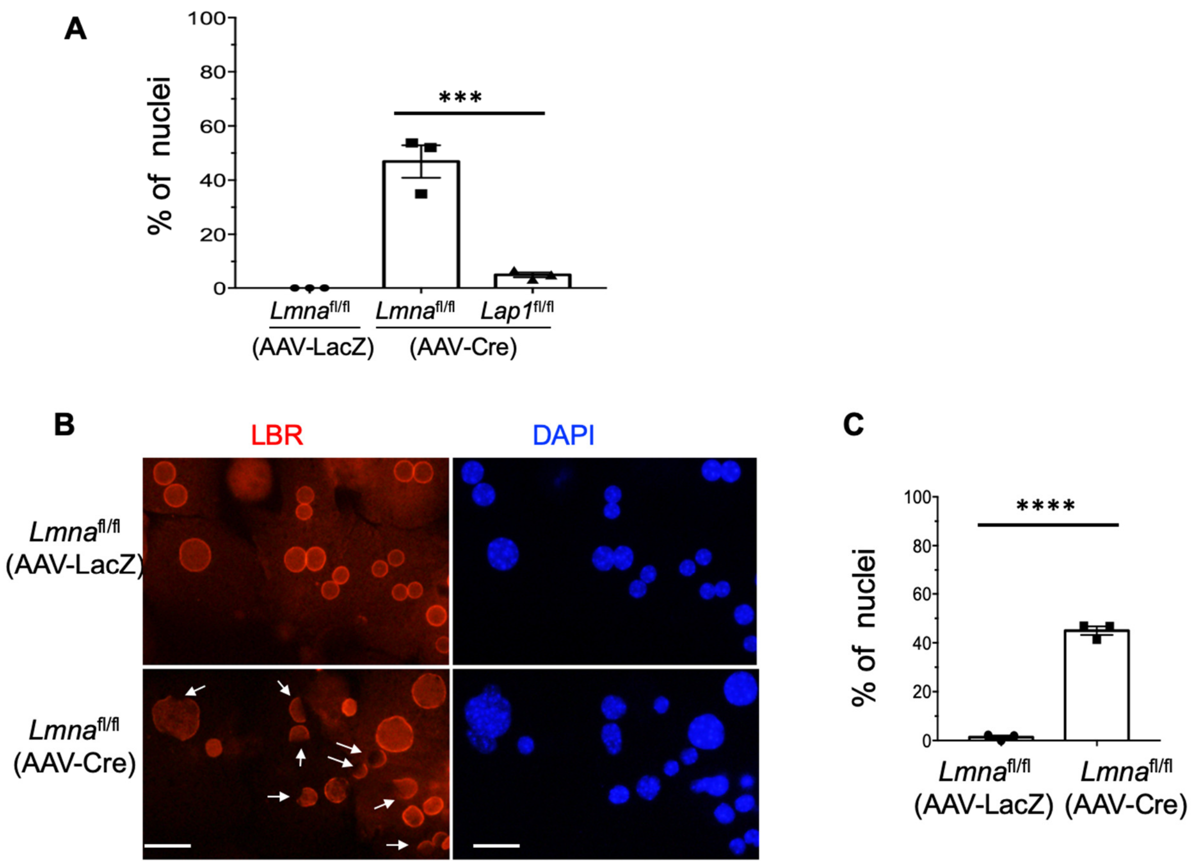


**Fig. S4.** (A) Mean percentages of hepatocytes with misshapen nuclei based on anti-lamin B1 Ab labeling shown in Fig. 6B. Columns show mean percentages of misshapen nuclei, black symbols indicate data from separate experiments (201-263 nuclei counted for each experiment and condition) and error bars show SEM. ***$P$ <0.001 by one-way ANOVA followed by Tukey's multiple comparison test. (B) Representative widefield fluorescence photomicrographs of hepatocytes from *Lmna*$^{fl/fl}$ mice isolated 4 weeks after AAV-LacZ or AAV-Cre virus injection, labeled with anti-LBR Abs (red) and DAPI (blue). Arrows indicate nuclei with absence of LBR in part of the nuclear envelope. Scale bars: 25 μm. (C) Mean percentages of misshapen hepatocyte nuclei based on anti-LBR Ab labeling. Columns show mean percentages of misshapen nuclei, black symbols indicate data from separate experiments (203-256 nuclei counted for each experiment and condition) and error bars show SEM. ****$P$ < 0.0001 by 2-tailed Student's $t$ test.

Supplemental Figure S5

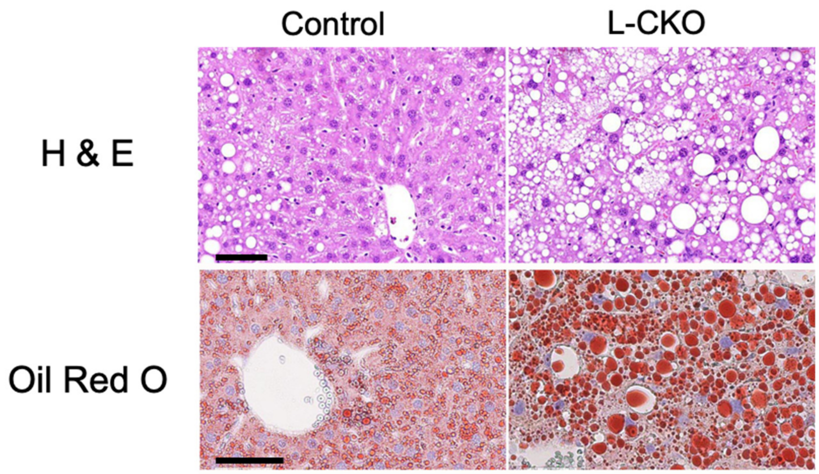


**Fig. S5**. Representative photomicrographs of liver sections from control (*Lap1*$^{fl/fl}$) and L-CKO mice at 3 months of age fed high fat diet for 8 weeks stained with H & E or Oil Red O. Scale bars: 50 µm.

Supplemental Figure S6

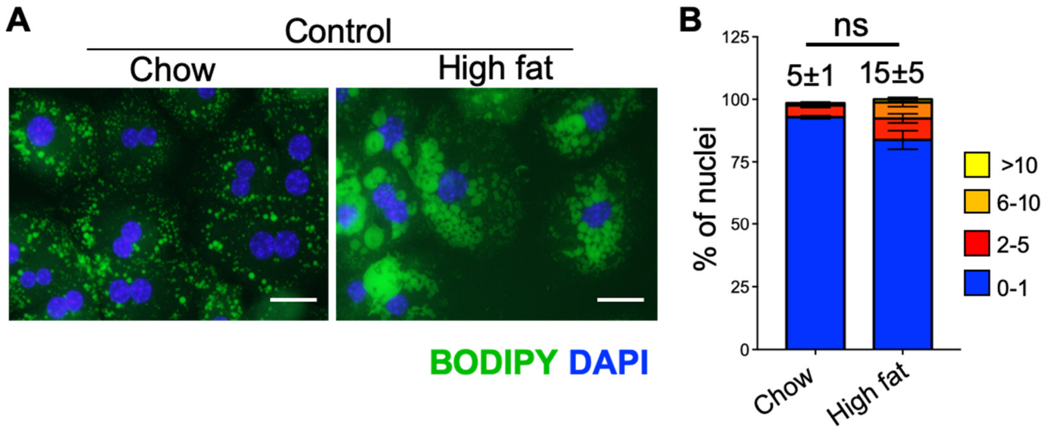


**Fig. S6.** (A) Representative widefield fluorescence photomicrographs of hepatocytes from control (*Lap1*$^{fl/fl}$) mice stained with BODIPY (green) and DAPI (blue). Hepatocytes were isolated from chow-fed (left panel) or high fat diet-fed (right panel) control mice. Scale bar: 25 µm. (B) Stacked column graph with different colors representing the percentages of hepatocyte nuclei containing the indicated numbers of nuclear LDs. We analyzed a total of 325 (chow diet) and 204 (high fat diet) nuclei of hepatocytes cultured on coverslips. The numbers at the top of the graphs indicate the mean percentages of hepatocyte nuclei with 2 or more nuclear LDs. These values and those within graphs are means ± SEM. ns = not significant by 2-tailed Student's *t* test.

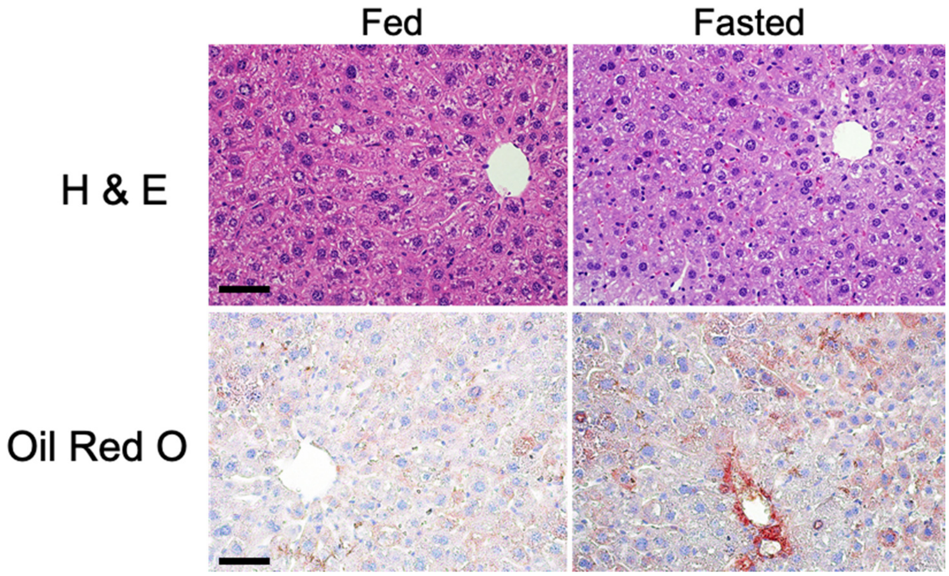


**Fig. S7**. Photomicrographs of liver sections from L-CKO mice at 3 months of age fed normally (Fed) or fasted for 24 hours (Fasted) stained with H & E or Oil Red O. Scale bars: 50 µm.

Supplemental Figure S8

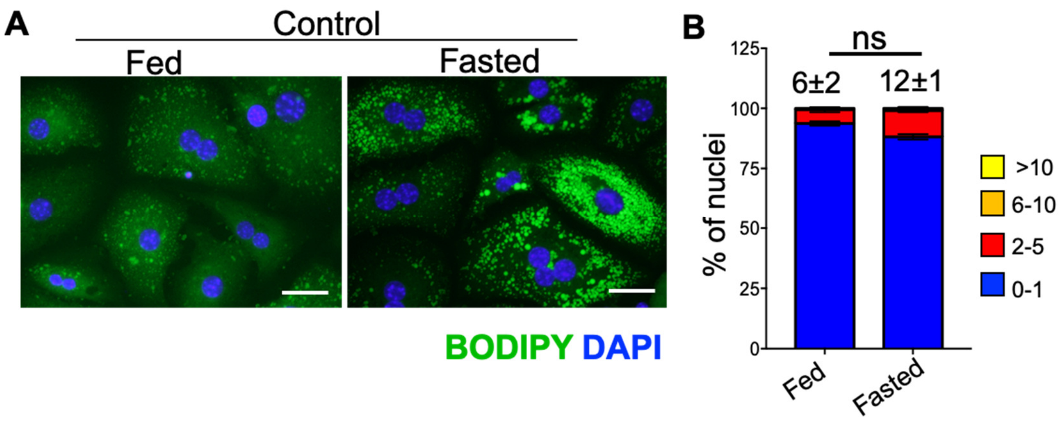


**Fig. S8.** (A) Representative widefield fluorescence photomicrographs of hepatocytes from control (*Lap1*$^{fl/fl}$) mice stained with BODIPY (green) and DAPI (blue). Hepatocytes were isolated from mice fed a normal chow diet (left panel) or mice fasted for 24 hours (right panel). Scale bar: 25 µm. (B) Stacked column graph with different colors representing the percentage of nuclei containing the indicated numbers of nuclear LDs in hepatocytes isolated from control mice fed normally or after 24 hours of fasting. We analyzed a total of 455 (fed) and 361 (fasted) nuclei of hepatocytes cultured on 3 different coverslips (n = 3 per group). The numbers at the top of the graphs indicate the mean percentages of hepatocyte nuclei with 2 or more nuclear LDs. These values and those within graphs are means ± SEM. ns = not significant by 2-tailed Student's *t* test.
